# Human *O*-linked Glycosylation Site Prediction Using Pretrained Protein Language Model

**DOI:** 10.1101/2023.10.23.563673

**Authors:** Subash C. Pakhrin, Neha Chauhan, Salman Khan, Jamie Upadhyaya, Charles Keller, Laurie N. Neuman, Moriah R. Beck, Eduardo Blanco

## Abstract

*O-*linked glycosylation of proteins is an essential post-translational modification process in *Homo sapiens*, where the attachment of a sugar moiety occurs at the oxygen atom of serine and/or threonine residues. This modification plays a pivotal role in various biological and cellular functions. While threonine or serine residues in a protein sequence are potential sites for *O-*linked glycosylation, not all threonine or serine residues are *O-*linked glycosylated. Furthermore, the modification is reversible. Hence, it is of vital importance to characterize if and when *O-*linked glycosylation occurs. We propose a multi-layer perceptron-based approach termed OglyPred-PLM which leverages the contextualized embeddings produced from the ProtT5-XL-UniRef50 protein language model that significantly improves the prediction performance of human *O-*linked glycosylation sites. OglyPred-PLM surpassed the performance of other indispensable *O-*linked glycosylation predictors on the independent benchmark dataset. This demonstrates that OglyPred-PLM is a powerful and unique computational tool to predict *O-*linked glycosylation sites in proteins and thus will accelerate the discovery of unknown *O-*linked glycosylation sites in proteins.

## Introduction

*O-*linked glycosylation is an important post-translational modification (PTM) in humans, involving the attachment of glycans to threonine (T) or serine (S) residues of protein sequences^1^. *O-*linked glycosylation sites can affect protein structure and function, resulting in various physiological and pathological outcomes, including protein dysfunction^2^ (namely, disruption in intercellular communication), cellular dysfunction, immune deficiencies^3^, hereditary disorders^4^, congenital disorders of glycosylation, and cancers^5,6^. Therefore, precise identification of *O-*linked glycosylation sites holds significant remedial potential in humans.

Experimental methods such as mass spectrometry^7,8^ are used to identify *O-*linked glycosylation sites. Through this technique, 9,354 *O-*glycosylation sites have been identified^9^. Even though this type of experiment is the most reliable method to identify *O-*linked glycosylation sites, there are several reasons that limit efficiency. Notably, these methods can be intricate, time-consuming, labor-intensive, and expensive. Therefore, the usage of statistical tools developed through machine learning (ML) and deep learning (DL) methodologies can become the prominent solution for characterizing *O-*linked glycosylation sites.

Recently there has been significant progress in ML and DL areas which has led to the creation of several computational tools that can predict *O-*linked glycosylation sites^10^. For example, NetOGlyc^11^ is a multilayer perceptron (MLP) based tool built for mucin-type *O-*linked glycosylation prediction. This approach used the accessible surface area and amino acid composition for encoding the amino acid sequences^12,13^. The Oglyc^14^ approach relies on the binary profile features and physicochemical properties of protein peptides. This method leverages support vector machine (SVM) algorithms to predict *O-* glycosylation sites. EnsembleGly^15^ uses ensemble SVM to predict *O-*glycosylation sites that utilize physicochemical properties, evolutionary features and intuitive binary encoding scheme to encode the protein peptide. The CKSAAP_OGlySite^16^ method uses SVM with the composition of k-spaced amino acids pairs (CKSAAP) feature encoding scheme to predict *O-*glycosylation sites. The GlycoPP^17^ tool utilized SVM to predict *O-*glycosylation sites where datasets were encoded with similar features used by EnsembleGly^15^ method. GlycoMine^18^ is a Random Forest (RF) based *O-*linked glycosylation site prediction and its datasets are numerically encoded with sequence and heterogeneous functional-based features. Positive unlabeled (PU)^19^ learning technique is used in GlycoMine_PU^20^ to detect *O-*linked glycosylation sites. SPRINT-Gly^21^ is a MLP based method which detects human and mouse *O-*glycosylation sites. Here, the protein peptide (length = 5) is encoded in a sequence and predicted secondary structural-based features. Captor^22^ is an *O-*glycosylation site prediction tool that uses the latest experimentally characterized OGP dataset. These datasets were encoded with sequence and physicochemical based features (Aaindex)^23^. These features were merged and utilized for training the SVM classifier. Recently, Alkuhlani et al.^24^ developed TAPE PLM-based^25^ *O-*linked glycosylation sites predictor and used XGBoost to classify the *O-* glycosylation site in the proteins.

The majority of the methods mentioned above^26-31^ have their input features manually curated for prediction. Furthermore, the *O-*linked glycosylation prediction method from Alkuhlani et al.^24^ is the only one that leveraged the benefits of embeddings from large protein language models (TAPE^25^). However, they have not extensively explored other complementary and recently developed PLMs. The *O-*linked glycosylation prediction method of Alkuhlani et al.^24^ still uses a window size of 31 around site of interest to extract the TAPE-based PLM features. Additionally, this method uses SVM based feature selection approaches which are indeed meticulous, yet also onerous. Moreover, a recent study^32^ evaluated the performance of different protein language models (PLM) for protein representation comparing them against each other. Through this study, it has been concluded that ProtT5-XL-Uniref50^33^ (herein called ProtT5) achieved the best performance in most of the proteomics tasks. However, ProtT5 has not yet been used for *O-*linked glycosylation site prediction. OglyPred-PLM utilizes high-quality experimentally characterized *O-*linked glycosylation data derived from the OGP database^9^, whose site of interest “S/T” is encoded by pre-trained PLM.

Recent developments in the field of natural language processing^34^ led to the creation of transformer-based large language models that have been trained on extensive and diverse corpora of unlabeled data. A wide range of PLMs have been developed^25,33,35,36^ which are trained with an enormous number of unlabeled protein sequences that are available in UniProt databases^37^ and other resources. Elnaggar et al. have developed ProtT5-XL-UniRef50^33^ PLM trained with 2.5 billion protein sequences where protein sequences are considered as sentences and individual amino acids as words. The representative embeddings produced from these models have been used for numerous posterior tasks. The results indicate that the feature embeddings generated by unsupervised PLM effectively encapsulate crucial characteristics encompassing the evolutionary context of a sequence, physicochemical properties, contact map, subcellular localization, distant interconnections in protein sequences, taxonomy, protein structure and function^38-43^. Likewise, features derived from these transformer-based pretrained PLMs have been proven to be effective in predicting a range of attributes, including intrinsic disorder sites^44^, signal peptides^45^, *N-*linked glycosylation sites^46^, subcellular localization^47^, phosphorylation sites^48^, protein structural featuers^49^, and binding residues^41^.

In this work, we developed a computation tool called OglyPred-PLM (*O-*linked glycosylation sites Prediction using pretrained Protein Language Model) that uses representative embeddings from a pre-trained ProtT5 PLM to train the MLP model and eventually strengthen the prediction performance of *O-* linked glycosylation sites. Similarly, we carry out a comprehensive comparison of the performance of ProtT5 PLM embeddings with other quintessential PLMs i.e., ESM2 (3B parameters)^35^ and ANKH^36^ approaches. The results demonstrate that the embeddings from ProtT5 are slightly better than other PLMs under consideration. Moreover, OglyPred-PLM was compared with the recent spectrum of *O-*linked glycosylation predictors. The experiments reveal that OglyPred-PLM was able to surpass the predictive performance of essential predictors like SPRINT-Gly, Captor, and Alkuhlani et al. *O-*linked glycosylation methods. OglyPred-PLM produces MCC, ACC, SN, PRE, and SP of 0.614, 0.807, 0.817, 0.801 and 0.797 respectively on Alkuhlani et al. independent test dataset. Similarly, it outperformed indispensable SPRINT-Gly and Captor *O-*linked glycosylation predictors by a wide margin. Hence, OglyPred-PLM is an important computational tool that can be used to detect *O-*linked glycosylation sites and accelerate discovery. All programs and data can be accessed via https://github.com/PakhrinLab/OglyPred-PLM

## Results and Discussion

### Generation of OGP Dataset using ProtT5 PLM

ProtT5 PLM was used to generate contextualized embeddings (dimensionality : 1024) for each amino acid site by using the full length of OGP protein sequences as an input. Only “S/T” embeddings were sent as an input to the OglyPred-PLM to train and predict *O-*linked glycosylation sites. We utilized an experimentally characterized OGP^9^ *O-*linked glycosylation dataset to train OglyPred-PLM. The psi-cd-hit^50^ tool (sequence identity threshold of 30 %) was applied to eliminate sequence redundancy from both the independent testing and training datasets, mitigating the risk of overfitting as well as maintaining diversity. Furthermore, we conducted a stratified 10-fold cross-validation grid search on the OGP training dataset to obtain optimal hyperparameters. Finally, we assessed the performance of the trained model using the independent test dataset and conducted a comparative analysis with established methods.

### Performance of models on The OGP Dataset

#### 10-fold Cross-Validation on the OGP Training Set with ProtT5 Features

To optimize hyperparameters^51^, we conducted a stratified 10-fold cross-validation (CV) on the OGP training dataset, and the results are presented in **Table 1**. Notably, the MLP architecture achieved superior results, with a mean MCC, mean ACC, mean SN, mean SP, and mean PRE of 0.664 ± 0.024, 0.831 ± 0.012, 0.833 ± 0.029, 0.830 ± 0.023, and 0.831 ± 0.016, respectively, for the 10-fold CV. The MLP architecture produced better results than other ML and DL models, thus, we determined MLP as our primary model and termed it OglyPred-PLM.

**Table 1.** Results of the 10-fold CV on the OGP training dataset using various models when training dataset are encoded with ProtT5 PLM. The most noteworthy values in each column have been emphasized in bold.

| Models | MCC ± 1 S.D. | ACC ± 1 S.D. | SN ± 1 S.D. | SP ± 1 S.D. | PRE ± 1 S.D. |
| --- | --- | --- | --- | --- | --- |
| LR | 0.623 ± 0.041 | 0.811 ± 0.020 | 0.814 ± 0.029 | 0.809 ± 0.014 | 0.809 ± 0.016 |
| XGBoost | 0.632 ± 0.023 | 0.816 ± 0.011 | 0.804 ± 0.013 | 0.827 ± 0.027 | 0.824 ± 0.022 |
| Random Forest | 0.602 ± 0.020 | 0.799 ± 0.010 | 0.738 ± 0.019 | **0.859 ± 0.017** | **0.840 ± 0.015** |
| SVM | 0.658 ± 0.017 | 0.829 ± 0.008 | 0.829 ± 0.015 | 0.829 ± 0.013 | 0.829 ± 0.011 |
| MLP | **0.664 ± 0.024** | **0.831 ± 0.012** | **0.833 ± 0.029** | 0.830 ± 0.023 | 0.831 ± 0.016 |
| 1D CNN | 0.651 ± 0.022 | 0.825 ± 0.011 | 0.819 ± 0.029 | 0.831 ± 0.030 | 0.830 ± 0.021 |

#### 10-fold Cross-Validation on the OGP Training Set with Ankh Features

We further explored the utility of the Ankh pre-trained PLM^36^ via 10-fold CV (type = stratified) on the OGP training dataset with Ankh PLM embeddings (**Table 2**). The MLP model used contextualized embeddings from Ankh PLM (feature vector length = 1,536) of the “S/T” token which produced impressive results with mean MCC, mean ACC, mean SN, mean SP, and mean PRE values of 0.658 ± 0.023, 0.828 ± 0.012, 0.831 ± 0.032, 0.825 ± 0.036, and 0.828 ± 0.025, respectively. These large, pre-trained PLMs exhibit an increased capacity to capture complex protein patterns, enhancing accuracy and generalization. However, 10-fold cross-validation with Ankh PLM slightly underperforms ProtT5 PLM embeddings. Hence, we chose pre-trained ProtT5 PLM to encode the protein sequence.

**Table 2.** Result of the 10-fold CV on the OGP training dataset using various models when training dataset are encoded with Ankh PLM. The most noteworthy values in each column have been emphasized in bold.

| Models | MCC ± 1 S.D. | ACC ± 1 S.D. | SN ± 1 S.D. | SP ± 1 S.D. | PRE ± 1 S.D. |
| --- | --- | --- | --- | --- | --- |
| LR | 0.614 ± 0.021 | 0.807 ± 0.010 | 0.791 ± 0.017 | 0.821 ± 0.010 | 0.816 ± 0.009 |
| XGBoost | 0.617 ± 0.024 | 0.808 ± 0.012 | 0.784 ± 0.020 | 0.833 ± 0.011 | 0.824 ± 0.011 |
| Random Forest | 0.563 ± 0.027 | 0.778 ± 0.014 | 0.696 ± 0.017 | **0.860 ± 0.014** | **0.832 ± 0.016** |
| SVM | 0.650 ± 0.035 | 0.825 ± 0.018 | 0.820 ± 0.018 | 0.830 ± 0.023 | 0.829 ± 0.021 |
| MLP | **0.658 ± 0.023** | **0.828 ± 0.012** | **0.831 ± 0.032** | 0.825 ± 0.036 | 0.828 ± 0.025 |
| 1D CNN | 0.625 ± 0.021 | 0.812 ± 0.011 | 0.818 ± 0.029 | 0.806 ± 0.031 | 0.809 ± 0.020 |

#### 10-fold Cross-Validation on the OGP Training Set with ESM2 (3B) Features

To encompass all PLM performance on *O-*linked glycosylation PTM prediction, the recently developed ESM2 (3B parameters) PLM^35^ last layer’s (layer no. 36) contextualized embedding was also taken into consideration. The feature vector size of ESM2 (3B) is 2,560 in length. The stratified 10-fold CV results are presented in **Table 3**. Surprisingly, the MLP architecture produced better results than other ML and DL methods across ProtT5, Ankh and ESM2 (3B) embeddings. The 10-fold CV result of ESM2 (3B) is slightly less than ProtT5 and ANKH PLM performance. Hence, we concluded that ProtT5 PLM’s embedding with MLP architecture is favorable for independent *O-*linked glycosylation PTM prediction purposes.

**Table 3.** Result of the 10-fold CV on the OGP training dataset using various models when training dataset are encoded with ESM2 (3B) PLM. The most noteworthy values in each column have been emphasized in bold.

| Models | MCC ± 1 S.D. | ACC ± 1 S.D. | SN ± 1 S.D. | SP ± 1 S.D. | PRE ± 1 S.D. |
| --- | --- | --- | --- | --- | --- |
| LR | 0.614 ± 0.034 | 0.806 ± 0.017 | 0.816 ± 0.015 | 0.797 ± 0.027 | 0.801 ± 0.022 |
| XGBoost | 0.577 ± 0.028 | 0.788 ± 0.014 | 0.766 ± 0.018 | 0.810 ± 0.022 | 0.802 ± 0.019 |
| Random Forest | 0.538 ± 0.028 | 0.763 ± 0.014 | 0.666 ± 0.026 | **0.861 ± 0.015** | **0.828 ± 0.016** |
| SVM | 0.616 ± 0.024 | 0.808 ± 0.012 | 0.832 ± 0.017 | 0.783 ± 0.020 | 0.793 ± 0.015 |
| MLP | **0.624 ± 0.037** | **0.811 ± 0.017** | **0.833 ± 0.029** | 0.788 ± 0.047 | 0.799 ± 0.032 |
| 1D CNN | 0.612 ± 0.025 | 0.805 ± 0.012 | 0.796 ± 0.025 | 0.814 ± 0.024 | 0.811 ± 0.017 |

### Testing on OGP Independent test Dataset with ProtT5 Features

To assess the performance, we trained OglyPred-PLM on the overall OGP training (controlled) dataset and evaluated it with an independent (uncontrolled) OGP *O-*glycosylation test dataset. All *O-*linked sites (and their respective protein sequences) used in OglyPred-PLM training are absent in the independent test dataset. Therefore, the model cannot learn representations from any *O-*linked sites in the independent test dataset. The model achieved MCC, ACC, PRE, SN, and SP values of 0.613, 0.806, 0.817, 0.789, and 0.824 (as shown in **Table 4**), respectively. The efficiency measures produced by OglyPred-PLM when the independent test dataset was not under-sampled are shown in **Supplementary Table S1**. Furthermore, OglyPred-PLM classified 308 samples as true negative (TN), 295 as true positive (TP), 66 samples as false positive (FP) and 79 as false negative (FN). Notably, OglyPred-PLM exhibited the highest AUC and PrAUC (**Figures** 1A and 1B) compared to other models, affirming its robustness for *O-*linked glycosylation site prediction.

**Figure 1.**
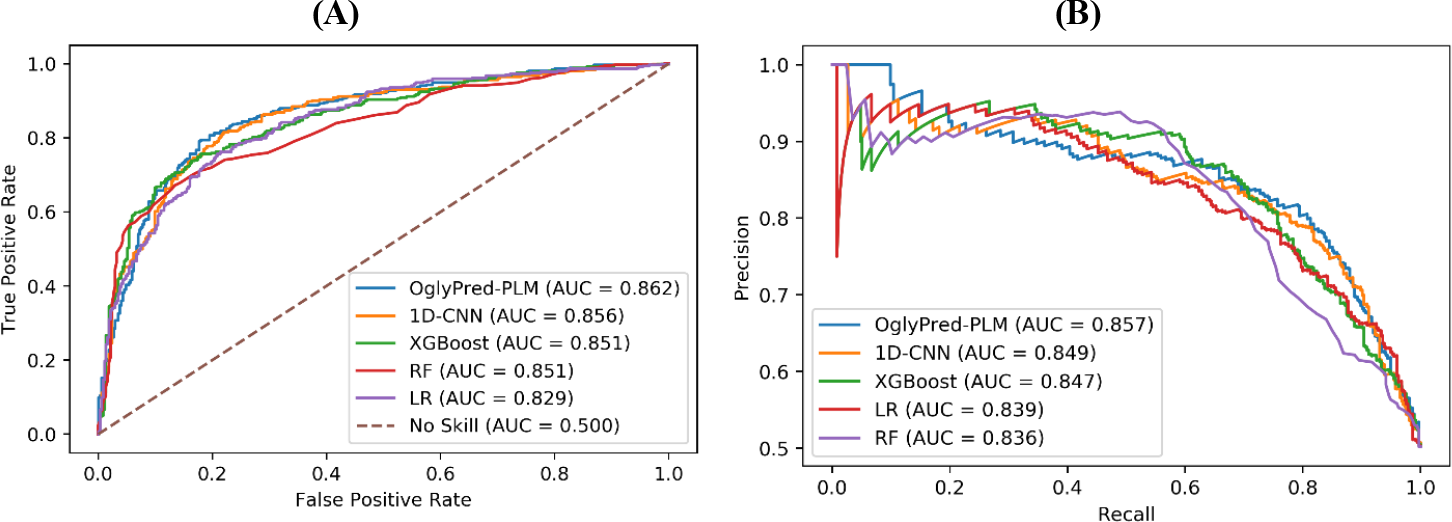
(A) Comparisons of ROC curves of OglyPred-PLM and other models on the OGP independent test dataset. Each model’s area under the ROC curve is reported. (B) Comparisons of precision-recall curves of OglyPred-PLM and other models on the OGP independent test dataset. Each model’s area under the PrAUC curve is reported.

**Table 4.**
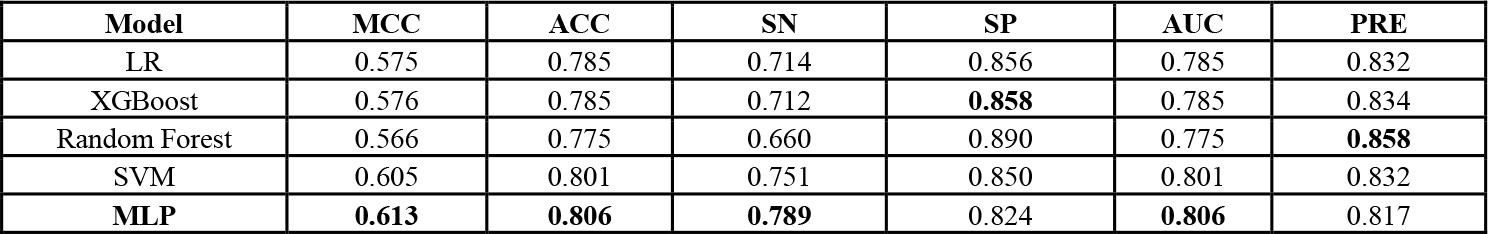
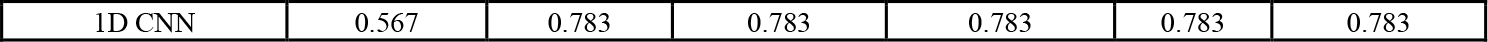
Performance metrics of various trained models on the OGP independent test set with ProtT5 features. The most noteworthy values in each column have been emphasized in bold.

| Model | MCC | ACC | SN | SP | AUC | PRE |
| --- | --- | --- | --- | --- | --- | --- |
| LR | 0.575 | 0.785 | 0.714 | 0.856 | 0.785 | 0.832 |
| XGBoost | 0.576 | 0.785 | 0.712 | **0.858** | 0.785 | 0.834 |
| Random Forest | 0.566 | 0.775 | 0.660 | 0.890 | 0.775 | **0.858** |
| SVM | 0.605 | 0.801 | 0.751 | 0.850 | 0.801 | 0.832 |
| MLP | **0.613** | **0.806** | **0.789** | 0.824 | **0.806** | 0.817 |
| 1D CNN | 0.567 | 0.783 | 0.783 | 0.783 | 0.783 | 0.783 |

### Testing on OGP Independent Test Dataset with Ankh Feature

The performance of the trained OglyPred-PLM, whose protein sequences were encoded by Ankh PLM, was evaluated with an OGP-independent test dataset. This trained model produced MCC, PRE, SN, SP, and ACC of 0.604, 0.810, 0.789, 0.816, and 0.802 respectively. The model classified the independent test dataset (unseen) samples as 305 TN, 295 TP, 69 FP, and 79 FN. **Table 5** shows independent test set results of various ML and DL models, whose protein sequences were encoded by ANKH-PLM.

**Table 5.** Performance metrics of various models on the OGP unseen test dataset with Ankh features. The most noteworthy values in each column have been emphasized in bold.

| Models | MCC | ACC | SN | SP | PRE |
| --- | --- | --- | --- | --- | --- |
| LR | 0.599 | 0.798 | 0.743 | 0.853 | 0.835 |
| XGBoost | 0.568 | 0.781 | 0.706 | 0.856 | 0.830 |
| Random Forest | 0.532 | 0.754 | 0.604 | **0.904** | **0.863** |
| SVM | 0.584 | 0.789 | 0.714 | 0.864 | 0.840 |
| MLP | **0.604** | **0.802** | **0.789** | 0.816 | 0.810 |
| 1D CNN | 0.591 | 0.795 | 0.789 | 0.802 | 0.799 |

To acknowledge the effects of under-sampling, **Supplementary Table S2** displays how various models, using Ankh features, performed when the independent test dataset was not under-sampled.

### Testing on OGP Independent Test Dataset with ESM2 (3B) Feature

The trained models with ESM2 (3B) PLM^35^ features were assessed with the independent test dataset. **Table 6** elaborates on the performance of the model with the independent test dataset. All the ML and DL models faired similarly. However, the results from ESM2 (3B) embeddings were still less than ProtT5 and ANKH PLM. The performance metrices, when the independent test dataset were not under sampled, are shown in **Supplementary Table S3**. Additionally, **Supplementary Table S4** shows the confusion matrix produced by OglyPred-PLM, also without under-sampling, using ProtT5, Ankh and ESM2 (3B) features. Furthermore, based upon MCC **Table 7** elaborates that OglyPred-PLM trained with ProtT5 PLM feature representation improves upon that of ANKH and ESM2 (3B) PLM.

**Table 6.** *Performance metrices of* various *models on the OGP unseen test dataset with ESM2 (3B) features. The most noteworthy values in each column have been emphasized in bold*.

| Models | MCC | ACC | SN | SP | PRE |
| --- | --- | --- | --- | --- | --- |
| LR | 0.586 | 0.793 | 0.791 | 0.795 | 0.794 |
| XGBoost | 0.586 | 0.790 | 0.720 | 0.860 | 0.837 |
| Random Forest | 0.535 | 0.754 | 0.599 | **0.909** | **0.869** |
| SVM | 0.600 | 0.799 | 0.770 | 0.829 | 0.818 |
| MLP | **0.602** | **0.801** | **0.795** | 0.807 | 0.805 |
| 1D CNN | 0.598 | 0.799 | 0.783 | 0.815 | 0.809 |

**Table 7.** Prediction performance of OglyPred-PLM with ProtT5, ANKH and ESM2 (B) features on the unseen test dataset. The most noteworthy values in each column have been emphasized in bold.

| PLM | MCC | ACC | SN | SP | PRE |
| --- | --- | --- | --- | --- | --- |
| ProtT5 | **0.613** | **0.806** | 0.789 | **0.824** | 0.806 |
| Ankh | 0.604 | 0.802 | 0.789 | 0.816 | **0.810** |
| ESM2 (3B) | 0.602 | 0.801 | 0.795 | 0.807 | **0.805** |

### Visualization Using t-distributed stochastic neighbor embedding Plot

The t-SNE^52^ method was used to discern the classification effectiveness of the embeddings from ProtT5 PLM and the second to last fully connected layer of the OglyPred-PLM model. This method projects these features into a two-dimensional space to identify class boundaries. The t-SNE’s learning rate was set to a value of 50 to visualize the scatter plot from ProtT5 features and thirty-two-dimensional feature vectors obtained from the second to last fully connected layer of the trained OglyPred-PLM framework. This learning rate was selected because it performed best among the range of learning rates considered, which was from 10 to 200 with a step size of 10. These plots were created using randomly selected set of 9,770 data points, evenly split between 4,885 negative and 4,885 positive samples.

*O-*linked glycosylated and non-*O-*linked glycosylated features were extracted from the ProtT5 PLM. **Figure 2 (A)** illustrates clusters of positive and negative samples extracted from ProtT5 PLM; however, the class boundaries of the positive and negative samples remain indistinct. **Figure 2 (B)** presents the scatter or the t-SNE plot of the features produced from the second to last fully connected layer of the trained OglyPred-PLM framework, where the positive samples (orange points) and negative samples (blue points) are distinctly clustered. **Figure 2 (B)** shows that the pre-trained per residue PLM feature embeddings from the site of interest, along with the trained MLP framework, is capable of learning *O-*linked glycosylation patterns and thus can classify negative and positive samples in two-dimensional space.

**Figure 2.**
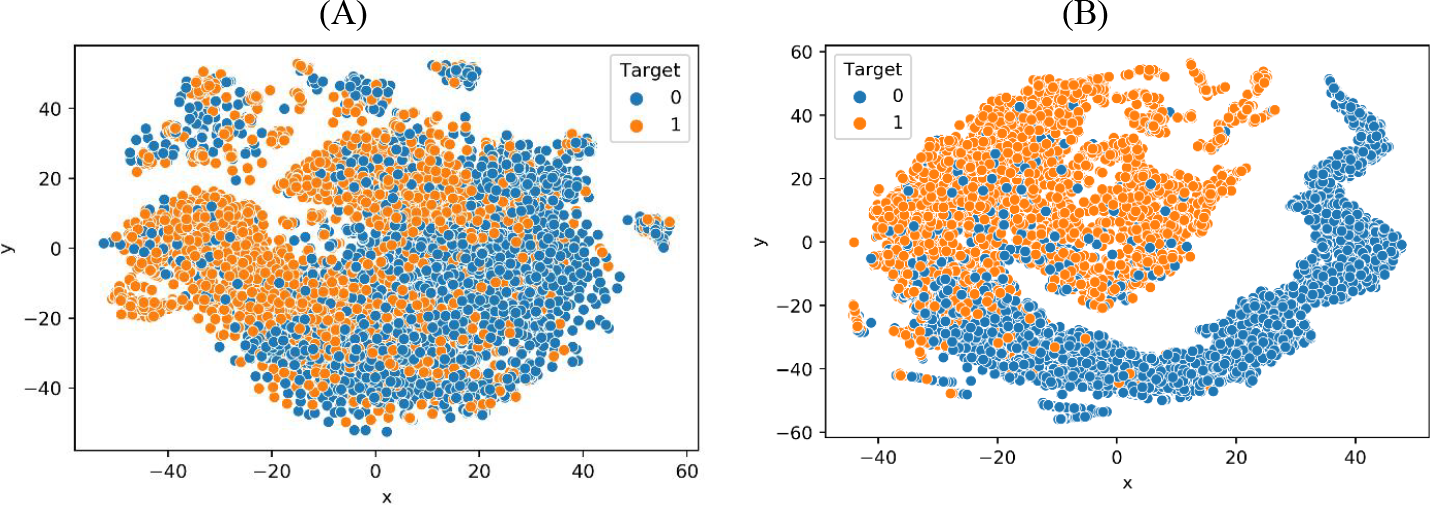
t-SNE visualization of the acquired features. (A) Features derived from the ProtT5 PLM, (B) learned features from the trained MLP model (OglyPred-PLM).

### Comparison of OglyPred-PLM with other *O-*linked Glycosylation Site Predictors

To assess the performance of OglyPred-PLM against other established predictors, we trained our model on the OGP training dataset which excludes both the independent test dataset and SPRINT-Gly independent test dataset. Moreover, to avoid possible bias, SPRINT-Gly^21^ used an under-sampling method to reduce the ratio of positive to negative samples to 1:3 for training. Hence, we also trained OglyPred-PLM with a 1:3 ratio. The trained model was evaluated without an under-sampled SPRINT-Gly independent test dataset. Our model produced MCC, ACC, SN, SP, and PRE values of 0.297, 0.857, 0.873, 0.856, and 0.124, respectively (as shown in **Table 8**). The results yielded by the model are better than the structural and sequential feature-based SPRING-Gly method, making the detection of *O-*linked glycosylation sites more efficient. It should be noted that OglyPred-PLM has remarkable positive *O*-linked glycosylation detection capability (87.3%) compared to SPRINT-GLY (32.9%). Additionally, the confusion matrix of the MLP architecture shows that the model detected 2,892 samples as TN and 69 as TP. However, it erroneously classified 484 samples as FP and 10 as FN. The results for the consensus-based model, GPP, GlycoMine, NetOGlyc and GlycoPP were adopted from SPRINT-Gly.

**Table 8.** Prediction performance of OglyPred-PLM compared to other existing O-linked glycosylation site predictors, evaluated on the SPRINT-Gly independent test set. The most noteworthy values in each column have been emphasized in bold.

| Methods | MCC | AUC | ACC | SN | SP | PRE |
| --- | --- | --- | --- | --- | --- | --- |
| OglyPred-PLM | **0.297** | **0.865** | 0.857 | **0.873** | 0.856 | 0.124 |
| SPRINT-Gly | 0.191 | 0.824 | 0.940 | 0.329 | 0.954 | **0.144** |
| Consensus-based model | 0.042 | 0.523 | 0.227 | 0.223 | 0.216 | 0.018 |
| GPP | 0.047 | 0.578 | 0.489 | 0.671 | 0.485 | 0.030 |
| GlycoMine | 0.100 | 0.667 | **0.978** | 0.027 | **0.999** | 0.400 |
| NetOGlyc | 0.128 | 0.749 | 0.633 | 0.785 | 0.629 | 0.047 |
| GlycoPP | 0.061 | 0.538 | 0.973 | 0.038 | 0.994 | 0.136 |

Moreover, to compare our method with other quintessential predictors like OGP and Captor, we removed the protein sequences from the OGP training dataset by excluding all *O-*linked and non-*O-*linked glycosylation sites that are present in the Captor independent test dataset and fitted the model with the resulting training dataset. We examined the trained model with the Captor independent test dataset. **Table 9** reveals that the OglyPred-PLM results are better than Captor and OGP predictors (superior positive *O*-linked glycosylation detection capability). OglyPred-PLM was successful at classifying 1,057 samples as TN and 261 samples as TP. However, it falsely classified 250 samples as FP and 79 samples as FN. It should be noted that OglyPred-PLM, Captor and OGP methods were calibrated and examined with the same experimentally verified OGP dataset.

**Table 9.**
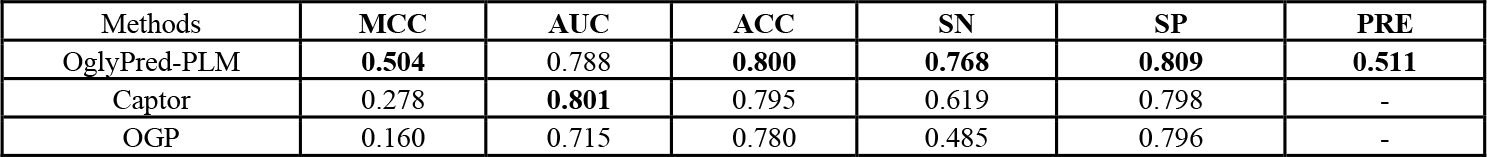
Prediction performance of OglyPred-PLM compared to other existing O-linked glycosylation site predictors on the Captor independent test set. The most noteworthy values in each column have been emphasized in bold.

Furthermore, to exhaustively compare OglyPred-PLM with PLM-based (TAPE) predictor we compared our results with this recently developed Alkuhlani et al. *O-*linked glycosylation method. Again, OglyPred-PLM shows improved results over their PLM-based method. These results are presented in **Table 10**.

**Table 10.** Prediction performance of OglyPred-PLM compared to Alkuhlani et al. PLM-based (TAPE) O-linked glycosylation site predictor on the Alkuhlani et al. independent test set. The most noteworthy values in each column have been emphasized in bold.

| Methods | MCC | AUC | ACC | SN | SP | PRE |
| --- | --- | --- | --- | --- | --- | --- |
| OglyPred-PLM | **0.614** | 0.807 | **0.807** | **0.817** | **0.797** | **0.801** |
| Alkuhlani et al. | 0.553 | **0.829** | 0.776 | 0.739 | 0.813 | - |

## Conclusion

In this work, an MLP-based model is used to detect *O-*linked glycosylation sites in amino acid sequences of proteins. OglyPred-PLM utilizes an embedding from pre-trained PLM (ProtT5) to encode amino acid sequences. The uniqueness of this approach is the use of contextualized embedding of the site of interest from a pre-trained PLM (ProtT5) and the use of a high-quality experimentally characterized *O-* linked glycosylation data set. The unseen test dataset results and comparison with other approaches show that OglyPred-PLM achieves more favorable performance over other approaches. Hence, it can be concluded that OglyPred-PLM is a trustworthy human *O-*linked glycosylation site prediction tool that exceeds prior tools in X.

The experiments demonstrate that the improvement in the performance of the OglyPred-PLM is due to following three factors: (i) the use of site of interest (“S/T”) embeddings extracted from the ProtT5 PLM contextualized file (ProtT5 PLM was fed with the entire protein .fasta file), (ii) training with a high-quality experimentally verified OGP database and (iii) the use of the well-trained and generalized MLP architecture. The t-SNE plot (**Figure 2** (B)) shows our trained model (OglyPred-PLM) can cluster *O-*linked and non-*O-*linked glycosylation serine or threonine residues in two-dimensional space. For any protein sequence, these ProtT5 PLMs embeddings can be easily extracted hence OglyPred-PLM provides fast and reliable predictions of *O-*linked glycosylation sites.

We foresee three avenues of future research to improve the performance of methods that utilize PLMs for *O-*linked and *N-*linked glycosylation PTM site prediction: (i) combining PLM features with other physicochemical features, (ii) using structural information predicted by AlphaFold2^53,54^ or ESMFold^55^ to build models that utilize graph networks^56^ and (iii) taking representative negative *O-*linked glycosylation sites from proteins dwelling within the Golgi apparatus subcellular localization of human cells. Finally, with the development of influential PLMs in the future, the prediction performance of the methods that make use of embeddings from these PLMs is more likely to improve.

## Methods

### Data Set

*O*-glycoprotein repository, known as OGP^9^, has the largest experimentally verified dataset for our research. It contains more than 9,354 *O-*linked glycosylation sites. We used this repository to extract 1,474 human *O-*linked glycoproteins to train and evaluate our models. Four proteins from the OGP repository were omitted due to their absence in the UniProt database. We used a psi-cd-hit^50^ tool to remove homologous glycoproteins that share 30 percent sequence similarity or more resulting in a subset of 1,176 glycoproteins. To avoid overestimation of performance, we made sure that no glycoproteins sequences from the independent test dataset were present in the training dataset as the PLM can learn its representation for other sites from the same glycoproteins^46^. Moreover, we split 1,176 glycoproteins into 1,059 training and 117 independent test glycoproteins. The annotated *O-*linked glycosylation sites from OGP database were defined as positive sites, and any remaining S or T sites from same glycoproteins were deemed as putative negative sites. **Table 11** presents the total number of positive and negative sites in the training and independent test datasets. To mitigate any biases toward the negative majority class, we balanced both the training and independent test dataset via under-sampling^57^.

**Table 11.** Positive and Negative O-linked Glycosylation Sites for Training and Independent Testing.

| Data set | Number of Proteins | Positive | Negative |
| --- | --- | --- | --- |
| Training | 1,059 | 4,885 | 114,144 |
| Test | 117 | 374 | 11,466 |

### Feature Encoding

An important process while developing a ML/DL model for the prediction of protein *O-*linked glycosylation sites is representing the primary amino acids with fixed size pertinent feature embeddings via a suitable encoding scheme^48,58^. We employed a pretrained PLM to generate per-residue contextualized embeddings for this investigation.

### Per-Residue Contextualized Embedding from ProtT5

The ProtT5-XL-UniRef50 (ProtT5) PLM was used to generate a contextualized per-residue embedding for target (“S/T”) sites. ProtT5 is a self-supervised learning model based on the T5 architecture^59^. The self-supervised model learns from the data itself, creating its own supervision signals without the need of external labels. Moreover, ProtT5 is trained with 2.5 billion unlabeled protein sequences from the BFD^60,61^ and Uni-Ref50^62^ databases. The entire protein sequence was fed into the pre-trained ProtT5 PLM to produce a contextualized per-residue feature embedding. Subsequently, site-of-interest (“S/T”) embeddings were extracted and fed into the MLP architecture.

### Model for Contextualized Embedding Obtained from ProtT5

The per-residue contextualized embedding produced from the last encoder layer of PLMs was used. The embeddings of the site of interest (“S/T”) were passed as an input to the MLP classification model. Furthermore, to select the appropriate embedding among ESM2 (3B parameters), ProtT5, and Ankh PLM, we performed a 10-fold CV on the training dataset. The 10-fold CV exhibits (as shown in the results section) that the ProtT5 PLM embeddings have superior protein representation compared to other PLMs considered. **Figure 3** illustrates the pipeline used in this work. Moreover, ProtT5, ESM2 (3B), and ANKH PLM use rotary positional embeddings^63^ and can generalize sequences longer than 1022 amino acids. The functionality of the OglyPred-PLM method is implemented by the MLP architecture which accepts ProtT5 representations of target sites (1024-length feature vectors) as input. The classifiers were constructed using Tensorflow^64^ as MLP models. Each model adopts an MLP architecture, consisting of three fully connected hidden layers with dimensions of 512, 256, and 32 respectively. Following each layer (except the final one), the rectified linear unit (ReLU) activation function is applied. To avoid over-fitting, a dropout rate of 0.3 was used after each fully connected layer. Moreover, techniques to avoid overfitting such as early stopping, ModelCheckpoint and reduce learning rate on plateau were used. The output layer uses sigmoid activation function with two neurons. This layer determines whether the given “S/T” amino acid is *O-*linked glycosylated (≥ 0.5) or non-*O-*linked glycosylated (< 0.5). A binary classifier was trained using the mini-batch training principle (with a batch size of 256) and Adam optimizer^65^ with a learning rate of 0.001. Since the datasets were imbalanced, the training and independent test datasets were loaded using random under-sampling. The MLP binary classifier was validated by assessing performance on the validation set after every training epoch and the model that achieved the highest MCC was selected. The trained model was then tested on the independent test (imbalanced or balanced) dataset, MCC and other binary classification metrics were reported. The hyperparameters used in OglyPred-PLM are shown in **Supplementary Table S5**.

**Figure 3.**
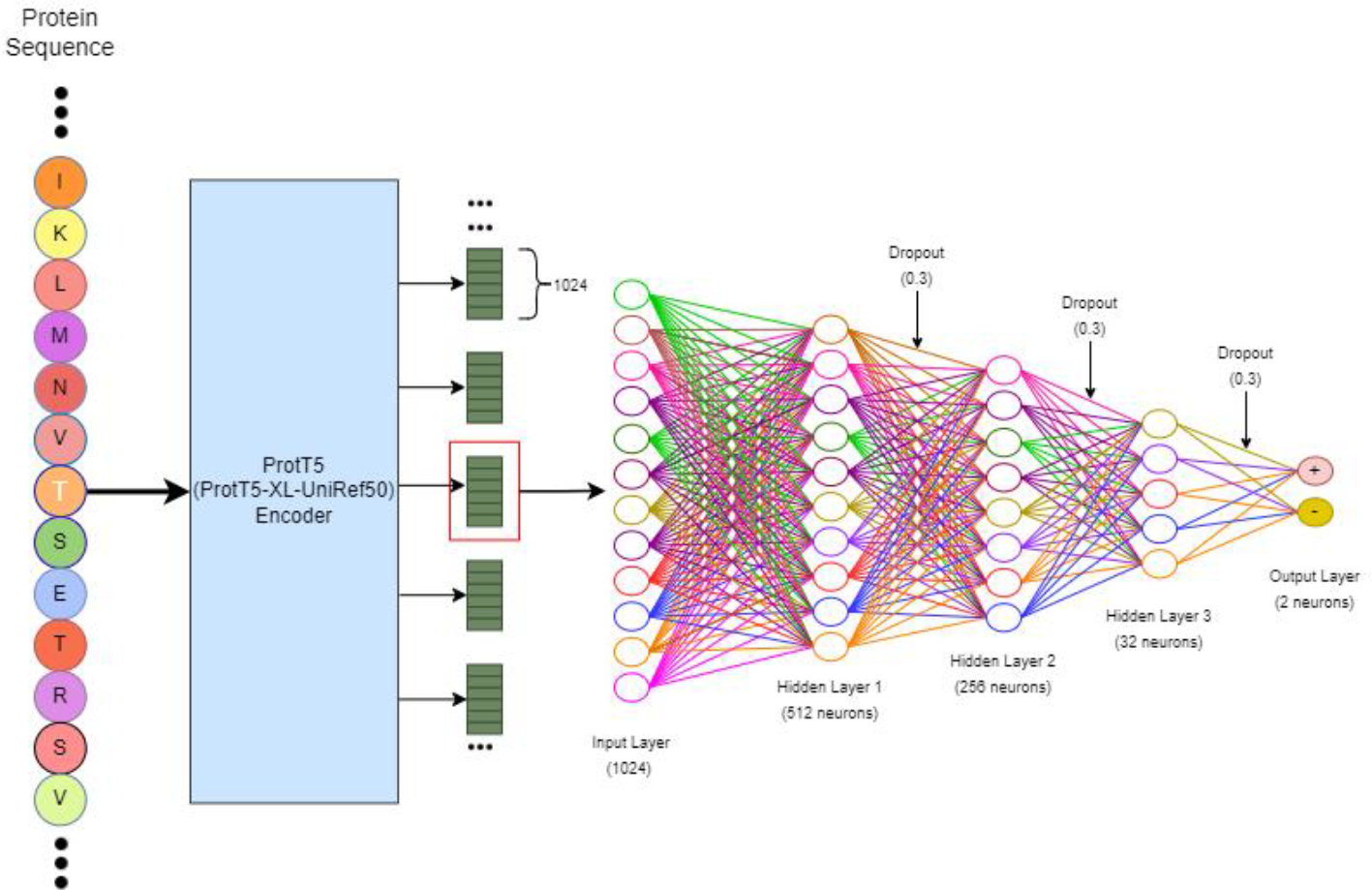
The overall framework of OglyPred-PLM. Beads with letters represent protein sequences. The sky-colored rectangular box represents ProtT5 PLM. Green rectangular boxes are 1024 per residue embeddings produced by ProtT5 PLM. Empty circles represent neurons. Each neuron is connected to other nodes via links like a biological axon synapse-dendrite connection^66^. A dropout of 0.3 means that 30 % of neurons are switched off randomly while training the MLP.

## Data availability

The developed tool, training and test data are available at https://github.com/PakhrinLab/OglyPred-PLM

## Acknowledgment

We acknowledge the use of the BeoShock and Beocat High-Performance Computing resources located at Wichita State University and Kansas State University. We appreciate the email discussions with Dr. Cangzhi Jia regarding their generated Captor dataset. In addition, we would also like to acknowledge the OGP dataset, ProtT5-XL-UniRef50, ESM2 and Ankh Protein Language Model made freely available to researchers.

## Author Contributions

S.C.P., M.R.B., and E.B. conceived and designed the experiments; S.C.P., N.C., and S.K. performed all the experiments and data analysis. J.A., L.N.N., and C.K. verified all the programs and models. S.C.P., N.C., S.K., J.A., L.N.N., C.K., M.R.B., E.B. edited and revised the manuscript. All authors have read and agreed to the published version of the manuscript.

## References

1 Yang, X.-m. in Advanced Research on Computer Education, Simulation and Modeling. (eds Song Lin & Xiong Huang) 445–450 (Springer Berlin Heidelberg).

2 Colley, K. J., Varki, A. & Kinoshita, T. in Essentials of Glycobiology (eds A. Varki et al.) 41–49 (2015).

3 Wolfert, M. A. & Boons, G. J. Adaptive immune activation: glycosylation does matter. Nat Chem Biol 9, 776–784, doi:10.1038/nchembio.1403 (2013).

4 Boskovski, M. T. et al. The heterotaxy gene GALNT11 glycosylates Notch to orchestrate cilia type and laterality. Nature 504, 456–459, doi:10.1038/nature12723 (2013).

5 Chen, Y., Zhou, W., Wang, H. & Yuan, Z. Prediction of O-glycosylation sites based on multi-scale composition of amino acids and feature selection. Med Biol Eng Comput 53, 535–544, doi:10.1007/s11517-015-1268-9 (2015).

6 Campos, D. et al. Probing the O-glycoproteome of gastric cancer cell lines for biomarker discovery. Mol Cell Proteomics 14, 1616–1629, doi:10.1074/mcp.M114.046862 (2015).

7 Agarwal, K. L., Kenner, G. W. & Sheppard, R. C. Feline gastrin. An example of peptide sequence analysis by mass spectrometry. J Am Chem Soc 91, 3096–3097, doi:10.1021/ja01039a051 (1969).

8 Medzihradszky, K. F. Peptide sequence analysis. Methods Enzymol 402, 209–244, doi:10.1016/S0076-6879(05)02007-0 (2005).

9 Huang, J. et al. OGP: A Repository of Experimentally Characterized O-glycoproteins to Facilitate Studies on O-glycosylation. Genomics Proteomics Bioinformatics 19, 611–618, doi:10.1016/j.gpb.2020.05.003 (2021).

10 KC, D. B. Computational Methods for Predicting Post-Translational Modification Sites. (Springer US, 2022).

11 Julenius, K., Mølgaard, A., Gupta, R. & Brunak, S. Prediction, conservation analysis, and structural characterization of mammalian mucin-type O-glycosylation sites. Glycobiology 15, 153–164, doi:10.1093/glycob/cwh151 (2005).

12 Klausen, M. S. et al. NetSurfP-2.0: Improved prediction of protein structural features by integrated deep learning. Proteins 87, 520–527, doi:10.1002/prot.25674 (2019).

13 Heffernan, R., Yang, Y., Paliwal, K. & Zhou, Y. Capturing non-local interactions by long short-term memory bidirectional recurrent neural networks for improving prediction of protein secondary structure, backbone angles, contact numbers and solvent accessibility. Bioinformatics 33, 2842–2849, doi:10.1093/bioinformatics/btx218 (2017).

14 Li, S., Liu, B., Zeng, R., Cai, Y. & Li, Y. Predicting O-glycosylation sites in mammalian proteins by using SVMs. Comput Biol Chem 30, 203–208, doi:10.1016/j.compbiolchem.2006.02.002 (2006).

15 Caragea, C., Sinapov, J., Silvescu, A., Dobbs, D. & Honavar, V. Glycosylation site prediction using ensembles of Support Vector Machine classifiers. BMC Bioinform. 8 (2007).

16 Chen, Y. Z., Tang, Y. R., Sheng, Z. Y. & Zhang, Z. Prediction of mucin-type O-glycosylation sites in mammalian proteins using the composition of k-spaced amino acid pairs. BMC Bioinformatics 9, 101, doi:10.1186/1471-2105-9-101 (2008).

17 Chauhan, J. S., Bhat, A. H., Raghava, G. P. & Rao, A. GlycoPP: a webserver for prediction of N- and O-glycosites in prokaryotic protein sequences. PLoS One 7, e40155, doi:10.1371/journal.pone.0040155 (2012).

18 Li, F. et al. GlycoMine: a machine learning-based approach for predicting N-, C- and O-linked glycosylation in the human proteome. Bioinformatics 31, 1411–1419, doi:10.1093/bioinformatics/btu852 (2015).

19 Bekker, J. & Davis, J. Learning from positive and unlabeled data: a survey. Machine Learning 109, 719–760, doi:10.1007/s10994-020-05877-5 (2020).

20 Li, F., Zhang, Y., Purcell, A. W. W., Geoffrey I. Chou, Kuo-Chen Lithgow, Trevor Li, C. & Song, J. Positive-unlabelled learning of glycosylation sites in thehuman proteome. BMC Bioinform. 20, 112 (2019).

21 Taherzadeh, G., Dehzangi, A., Golchin, M., Zhou, Y. & Campbell, M. P. SPRINT-Gly: Predicting N- and O-linked glycosylation sites of human and mouse proteins by using sequence and predicted structural properties. Bioinformatics 4140–4146. (2019).

22 Zhu, Y., Yin, S., Zheng, J., Shi, Y. & Jia, C. O-glycosylation site prediction for Homo sapiens by combining properties and sequence features with support vector machine. J Bioinform Comput Biol 20, 2150029, doi:10.1142/s0219720021500293 (2022).

23 Kawashima, S. et al. AAindex: amino acid index database, progress report 2008. Nucleic Acids Res 36, D202–205, doi:10.1093/nar/gkm998 (2008).

24 Alkuhlani, A., Gad, W., Roushdy, M. & Salem, A.-B. Prediction Of O-Glycosylation Site Using Pre-Trained Language Model And Machine Learning. International Journal of Intelligent Computing and Information Sciences 23, 41–52, doi:10.21608/ijicis.2023.160986.1218 (2023).

25 Rao, R. B. Nicholas et al. in Adv Neural Inf Process Syst (2019).

26 Hamby, S. E. & Hirst, J. D. Prediction of glycosylation sites using random forests. BMC Bioinformatics 9, 500, doi:10.1186/1471-2105-9-500 (2008).

27 Li, F. et al. GlycoMine(struct): a new bioinformatics tool for highly accurate mapping of the human N-linked and O-linked glycoproteomes by incorporating structural features. Sci Rep 6, 34595, doi:10.1038/srep34595 (2016).

28 Pakhrin, S. C., Aoki-Kinoshita, K. F., Caragea, D. & Kc, D. B. DeepNGlyPred: A Deep Neural Network-Based Approach for Human N-Linked Glycosylation Site Prediction. Molecules 26, 7314, doi:10.3390/molecules26237314 (2021).

29 Dhakal, A., Gyawali, R., Wang, L. & Cheng, J. A large expert-curated cryo-EM image dataset for machine learning protein particle picking. Scientific Data 10, 392, doi:10.1038/s41597-023-02280-2 (2023).

30 Pakhrin, S. C., Pokharel, S., Saigo, H. & Kc, D. B. Deep Learning-Based Advances In Protein Posttranslational Modification Site and Protein Cleavage Prediction. 2022/06/14 edn, Vol. 2499 (2022).

31 Pakhrin, S. C., Shrestha, B., Adhikari, B. & Kc, D. B. Deep Learning-Based Advances in Protein Structure Prediction. Int J Mol Sci 22, doi:10.3390/ijms22115553 (2021).

32 Unsal, S. et al. Learning functional properties of proteins with language models. Nature Machine Intelligence 4, 227–245, doi:10.1038/s42256-022-00457-9 (2022).

33 Elnaggar, A. et al. ProtTrans: Towards Cracking the Language of Lifes Code Through Self-Supervised Deep Learning and High Performance Computing. IEEE Trans Pattern Anal Mach Intell PP, doi:10.1109/TPAMI.2021.3095381 (2021).

34 Vaswani, A. e. a. Attention is all you need. In Proceedings of 31st International Conference on Neural Information Processing Systems (NIPS 2017) 1, 6000–6010 (2017).

35 Lin, Z. et al. Evolutionary-scale prediction of atomic-level protein structure with a language model. Science 379, 1123–1130, doi:10.1126/science.ade2574 (2023).

36 Elnaggar, A. et al. Ankh ☥: Optimized Protein Language Model Unlocks General-Purpose Modelling. bioRxiv, 2023.2001.2016.524265, doi:10.1101/2023.01.16.524265 (2023).

37 UniProt, C. UniProt: the universal protein knowledgebase in 2021. Nucleic Acids Res 49, D480–D489, doi:10.1093/nar/gkaa1100 (2021).

38 Pakhrin, S. C. Deep learning-based approaches for prediction of post-translational modification sites in proteins, Wichita State University, (2022).

39 Weissenow, K., Heinzinger, M. & Rost, B. Protein language-model embeddings for fast, accurate, and alignment-free protein structure prediction. Structure 30, 1169–1177 e1164, doi:10.1016/j.str.2022.05.001 (2022).

40 Nallapareddy, V. et al. CATHe: Detection of remote homologues for CATH superfamilies using embeddings from protein language models. bioRxiv, doi:10.1101/2022.03.10.483805 (2022).

41 Littmann, M., Heinzinger, M. & Dallago, C. Protein embeddings and deep learning predict binding residues for various ligand classes. Scientific reports 11, doi:10.1038/s41598-021-03431-4 (2021).

42 Zhang, S. et al. Applications of transformer-based language models in bioinformatics: a survey. Bioinformatics Advances 3, doi:10.1093/bioadv/vbad001 (2023).

43 Heinzinger, M. et al. Contrastive learning on protein embeddings enlightens midnight zone. NAR Genom Bioinform 4, lqac043, doi:10.1093/nargab/lqac043 (2022).

44 Song, Y. et al. Fast and accurate protein intrinsic disorder prediction by using a pretrained language model. Brief Bioinform, doi:10.1093/bib/bbad173 (2023).

45 Teufel, F. et al. SignalP 6.0 predicts all five types of signal peptides using protein language models. Nat Biotechnol 40, 1023–1025 doi:10.1038/s41587-021-01156-3 (2022).

46 Pakhrin, S. C. et al. LMNglyPred: prediction of human N-linked glycosylation sites using embeddings from a pre-trained protein language model. Glycobiology, doi:10.1093/glycob/cwad033 (2023).

47 Thumuluri, V., Almagro Armenteros, J. J., Johansen, A. R., Nielsen, H. & Winther, O. DeepLoc 2.0: multi-label subcellular localization prediction using protein language models. Nucleic Acids Res 50, W228–W234, doi:10.1093/nar/gkac278 (2022).

48 Pakhrin, S. C. et al. LMPhosSite: A Deep Learning-Based Approach for General Protein Phosphorylation Site Prediction Using Embeddings from the Local Window Sequence and Pretrained Protein Language Model. J Proteome Res 22, 2548–2557, doi:10.1021/acs.jproteome.2c00667 (2023).

49 Høie, M. H. et al. NetSurfP-3.0: accurate and fast prediction of protein structural features by protein language models and deep learning. Nucleic Acids Res 50, W510–W515, doi:10.1093/nar/gkac439 (2022).

50 Huang, Y., Niu, B., Gao, Y., Fu, L. & Li, W. CD-HIT Suite: a web server for clustering and comparing biological sequences. Bioinformatics 26, 680–682, doi:10.1093/bioinformatics/btq003 (2010).

51 Yang, L. & Shami, A. On hyperparameter optimization of machine learning algorithms: Theory and practice. Neurocomputing 415, 295–316, doi:10.1016/j.neucom.2020.07.061 (2020).

52 Maaten, L. v. d. & Hinton, G. Visualizing Data using t-SNE. Mach. Learn. Res. 9, 2579–2605 (2008).

53 Jumper, J. et al. Highly accurate protein structure prediction with AlphaFold. Nature 596, 583–589, doi:10.1038/s41586-021-03819-2 (2021).

54 Yuan, Q. et al. AlphaFold2-aware protein-DNA binding site prediction using graph transformer. Brief Bioinform 23, doi:10.1093/bib/bbab564 (2022).

55 Lin, Z. et al. Language models of protein sequences at the scale of evolution enable accurate structure prediction. bioRxiv, 2022.2007.2020.500902, doi:10.1101/2022.07.20.500902 (2022).

56 Yuan, Q., Chen, J., Zhao, H., Zhou, Y. & Yang, Y. Structure-aware protein-protein interaction site prediction using deep graph convolutional network. Bioinformatics 38, 125–132, doi:10.1093/bioinformatics/btab643 (2021).

57 Lemaitre, G., Nogueira, F. & Aridas, C. K. Imbalanced-learn: A Python Toolbox to Tackle the Curse of Imbalanced Datasets in Machine Learning. J. Mach. Learn. Res. 18, 559–563 (2017).

58 Y. Xu, Y.-X. D., J. Ding, Y.-H. Lei, L.-Y. Wu, N.-Y. Deng. iSuc-PseAAC: predicting lysine succinylation in proteins by incorporating peptide position-specific propensity. Sci. Rep. 5, 10184 (2015).

59 Raffel, C. et al. Exploring the limits of transfer learning with a unified text-to-text transformer. J. Mach. Learn. Res. 21, 1–67 (2020).

60 Steinegger, M., Mirdita, M. & Soding, J. Protein-level assembly increases protein sequence recovery from metagenomic samples manyfold. Nat Methods 16, 603–606, doi:10.1038/s41592-019-0437-4 (2019).

61 Steinegger, M. & Söding, J. Clustering huge protein sequence sets in linear time. Nature Communications 9, 2542, doi:10.1038/s41467-018-04964-5 (2018).

62 Suzek, B. E., Wang, Y., Huang, H., McGarvey, P. B. & Wu, C. H. UniRef clusters: a comprehensive and scalable alternative for improving sequence similarity searches. Bioinformatics 31, 926–932, doi:10.1093/bioinformatics/btu739 (2015).

63 Su, J. et al. ROFORMER: ENHANCED TRANSFORMER WITH ROTARY POSITION EMBEDDING. arXiv (2022).

64 Abadi, M. et al. Tensorflow: A System for Large-Scale Machine Learning. 12th Symposium on Operating Systems Design and Implementation, 265–283 (2016).

65 Kingma, D. P. B. J. Adam: A Method for Stochastic Optimization. arXiv e-prints, doi:https://ui.adsabs.harvard.edu/abs/2014arXiv1412.6980K (2014).

66 Südhof, T. C. The cell biology of synapse formation. J Cell Biol 220, doi:10.1083/jcb.202103052 (2021).

