## Supplemental Table 1-6 for "Human *O*-linked Glycosylation Site Prediction Using Pretrained Protein Language Model"

### Supplementary Materials:

*Table S1. Performance metrics of various trained models on the OGP independent test set (without under-sampling) with ProtT5 features. The most noteworthy values in each column have been emphasized in bold.*

| Models | MCC | ACC | SN | SP | AUC | PRE |
|---|---|---|---|---|---|---|
| LR | 0.2492 | 0.8285 | 0.7219 | 0.8320 | 0.7769 | 0.1229 |
| XGBoost | 0.2739 | 0.8516 | 0.7219 | 0.8558 | 0.7887 | 0.1404 |
| RF | 0.2418 | 0.8173 | 0.7299 | 0.8201 | 0.7750 | 0.1169 |
| SVM | 0.2651 | 0.8438 | 0.7219 | 0.8478 | 0.7848 | 0.1339 |
| **MLP** | **0.2744** | **0.8260** | **0.7914** | **0.8271** | **0.8098** | **0.1299** |
| 1D CNN | 0.2647 | 0.8342 | 0.7459 | 0.8371 | 0.7915 | 0.1300 |

*Table S2. Performance metrics of various trained models on the OGP independent test set (without under-sampling) with Ankh features. The most noteworthy values in each column have been emphasized in bold.*

| Models | MCC | ACC | SN | SP | AUC | PRE |
|---|---|---|---|---|---|---|
| LR | 0.2701 | 0.8521 | 0.7112 | 0.8567 | 0.7839 | 0.1393 |
| XGBoost | 0.2645 | 0.8423 | 0.7245 | 0.8461 | 0.7853 | 0.1331 |
| RF | 0.2529 | 0.8744 | 0.6069 | 0.8831 | 0.7450 | 0.1448 |
| SVM | 0.2610 | 0.8480 | 0.7005 | 0.8528 | 0.7767 | 0.1344 |
| **MLP** | **0.2719** | **0.8574** | **0.7005** | **0.8625** | **0.7815** | **0.1425** |
| 1D CNN | 0.2653 | 0.8419 | 0.7272 | 0.8457 | 0.7864 | 0.1332 |

*Table S3. Performance metrics of various trained models on the OGP independent test set (without under-sampling) with ESM2 (3B) features. The most noteworthy values in each column have been emphasized in bold.*

| Models | MCC | ACC | SN | SP | AUC | PRE |
|---|---|---|---|---|---|---|
| LR | 0.2345 | 0.7764 | 0.7639 | 0.7768 | 0.7704 | 0.1128 |
| XGBoost | 0.2630 | 0.8355 | 0.7018 | 0.8405 | 0.7712 | 0.1405 |
| RF | 0.2681 | 0.8714 | 0.6149 | 0.8809 | 0.7479 | 0.1609 |
| SVM | 0.2665 | 0.7944 | 0.8074 | 0.7939 | 0.8007 | 0.1270 |
| **MLP** | **0.2718** | **0.8340** | **0.7267** | **0.8380** | 0.7823 | **0.1428** |
| 1D CNN | 0.2672 | 0.8382 | 0.7049 | 0.8432 | 0.7740 | 0.1431 |

*Table S4. confusion matrix produced by OglyPred-PLM without under sample OGP independent test dataset using ProtT5, Ankh and ESM (3B) features.*

| | PLM | TN | FP | FN | TP |
|---|---|---|---|---|---|
| S/T Sites | ProtT5 | 9,484 | 1,982 | 78 | 296 |
| | Ankh | 9,890 | 1,576 | 112 | 262 |
| | ESM2 (3B) | 7,264 | 1,404 | 88 | 234 |

**Machine Learning and Deep Learning Models**

We briefly describe the machine learning and deep learning models used in this study.

Support Vector Machine[1] is a class of supervised machine learning algorithms used for classification and regression tasks[2]. The basic idea behind SVM is to find an optimal hyperplane that separates the data into different classes. When the data is not linearly separable, SVM can still classify it by using the kernel trick. The kernel trick maps the input data into a higher-dimensional feature space[3], where it might become linearly separable.

Random Forest[4] is a popular ensemble learning method used for classification and regression tasks in ML[4]. It is an extension of decision trees and combines multiple decision trees to make predictions. For classification tasks, it predicts the class label by taking a majority vote among the individual trees. Each tree's prediction is counted, and the class with the most votes becomes the final prediction.

Logistic Regression is an ML algorithm used for binary classification tasks[5]. It predicts the probability of an instance belonging to a certain class by fitting a logistic (sigmoid) function to the input features. It estimates coefficients to create a linear decision boundary that separates the two classes. XGBoost (Extreme Gradient Boosting) belongs to the family of gradient boosting methods. It sequentially adds weak models (decision trees) to iteratively correct the errors made by previous models. It optimizes a specific loss function by finding the best-fitting model in an additive manner.

XGBoost[6] (Extreme Gradient Boosting) is an ML classification and regression algorithm that belongs to the ensemble-based learning family. It possesses high predictive performance and efficiency.

1D Convolutional Neural Network[7] is a variant of convolutional neural networks (CNNs) specifically designed for processing one-dimensional sequential data[8]. It utilizes one-dimensional convolutional filters to capture local patterns and features in sequential data. The filters slide along the input sequence, performing convolutions and generating feature maps. While traditional CNNs are commonly used for image analysis and computer vision, 1D CNN works particularly well for sequential data. The hyperparameters and other details are explained in Supplementary Table S6.

*Table S5. Hyperparameters of OglyPred-PLM.*

| Name of the Parameters | Value Used |
|---|---|
| No. of layers | 4 |
| No. neuron in the first hidden layers | 512 |
| No. neuron in the second hidden layers | 256 |
| No. neuron in the third hidden layers | 32 |
| No. of neuron in the output layer | 2 |
| Activation Function | ReLU |
| Activation Function at output layer | SoftMax |
| Optimizer | Adam |
| Learning rate | 0.001 |
| Loss function | Binary cross entropy |
| Model Checkpoint | Monitor = 'Validation accuracy' |
| Reduce learning rate on plateau | Factor = 0.001 |
| Early stopping | Patience = 5 |
| Dropout | 0.3 |
| Decision Boundary | 0.5 |
| Batch size | 256 |
| Epochs | 400 |

*Table S6. Hyperparameters of ProtT5 features-based machine learning models.*

| Model name | Hyperparameters |
|---|---|
| Random Forest | n_estimators = 100, criterion = entropy |
| XGBoost | max_depth = 3, subsample = 0.8, n_estimators = 200, learning_rate = 0.05, random_state = 5 |
| SVM | C = 0.1, Probability = True, kernel='rbf' |
| Logistic Regression | C = 0.01 |
| 1D Convolutional Neural Network | filters=64, kernel_size=3<br>MaxPooling1D pool_size=2 |

### Model evaluation and performance metrics

In this study, non-*O*-linked glycosylation sites are considered negative sites, *and O*-linked glycosylation sites are considered positive sites. Non-*O*-linked glycosylation sites and *O*-linked glycosylation sites predicted correctly by the models are true negative (TN) and true positive (TP), respectively. The positive sites misclassified as negative sites are false negative (FN) and negative sites misclassified as positive sites are false positive (FP). A ten-fold cross-validation grid search was utilized to optimize the hyperparameters of the model. In ten-fold cross-validation, the training data is divided into ten equal parts. Nine parts of the divided training dataset are used to train the model and the remaining part is used for validation purposes. This mechanism iterated ten times. The results of 10-fold cross-validation, metrics are reported as the mean value ± one standard deviation.

Matthew's correlation coefficient (MCC), precision (PRE), accuracy (ACC), sensitivity (SN), and specificity (SP) were measured to observe the model performance. MCC shows the predictive capability of the trained model in a balanced fashion by considering all the elements of the confusion matrix (Equation (1)). Precision describes how many of the predicted positive cases are true positive (Equation (2)). Accuracy measures the overall correctness of a classification model's predictions (Equation (3)). When the negative and positive classes are highly imbalanced ACC may not be a trustworthy metric. Hence, random under-sampling technique is used to balance the dataset. Meanwhile, SN defines the model's ability to correctly identify positive residues (Equation (4)). On the other hand, SP measures the model's ability to correctly identify the negative residues (Equation (5)). Furthermore, the area under the precision-recall curve (PrAUC) and the area under the receiver operating characteristics (ROC) curve were also used as performance metrics. Importantly, it should be noted that a greater ROC curve signifies an improved classification model.

$$MCC = \frac{(TP)(TN)-(FP)(FN)}{\sqrt{(TP+FP)(TP+FN)(TN+FP)(TN+FN)}} \tag{1}$$

$$Precision = \frac{TP}{TP+FP} \times 100 \tag{2}$$

$$Accuracy = \frac{TP+TN}{TP+TN+FP+FN} \times 100 \tag{3}$$

$$Sensitivity = \frac{TP}{TP+FN} \times 100 \tag{4}$$

$$Specificity = \frac{TN}{TN+FP} \times 100 \tag{5}$$

### References


1 Cortes, C. & Vapnik, V. Support-vector networks. *Machine Learning* **20**, 273-297, doi:10.1007/BF00994018 (1995).

2 Smola, A. J. & Schölkopf, B. A tutorial on support vector regression. *Statistics and Computing* **14**, 199-222, doi:10.1023/B:STCO.0000035301.49549.88 (2004).

3 Zhao, Z.-j. & Jiao, M.-y. in *Proceedings of the 9th International Conference on Advanced Intelligent Systems and Informatics 2023.* (eds AboulElla Hassanien *et al.*) 521-531 (Springer Nature Switzerland).

4 Breiman, L. Random Forests. *Machine Learning* **45**, 5-32, doi:10.1023/A:1010933404324 (2001).

5 Sperandei, S. Understanding logistic regression analysis. *Biochem Med (Zagreb)* **24**, 12-18, doi:10.11613/bm.2014.003 (2014).

6 Chen, T. & Guestrin, C. in *Proceedings of the 22nd ACM SIGKDD International Conference on Knowledge Discovery and Data Mining* 785–794 (Association for Computing Machinery, San Francisco, California, USA, 2016).

7 Lecun, Y., Bottou, L., Bengio, Y. & Haffner, P. Gradient-based learning applied to document recognition. *Proceedings of the IEEE* **86**, 2278-2324, doi:10.1109/5.726791 (1998).

8 LeCun, Y., Bengio, Y. & Hinton, G. Deep learning. *Nature* **521**, 436-444, doi:10.1038/nature14539 (2015).
